# Single worm long read sequencing reveals genome diversity in free-living nematodes

**DOI:** 10.1101/2023.04.17.537128

**Authors:** Yi-Chien Lee, Hsin-Han Lee, Huei-Mien Ke, Yu-Ching Liu, Min-Chen Wang, Yung-Che Tseng, Taisei Kikuchi, Isheng Jason Tsai

## Abstract

Obtaining sufficient genetic material from a limited biological source is currently the primary operational bottleneck in studies investigating biodiversity and genome evolution. In this study, we employed multiple displacement amplification (MDA) and Smartseq2 to amplify nanograms of genomic DNA and mRNA, respectively from individual *Caenorhabditis elegans*. Although reduced genome coverage was observed in repetitive regions, we produced assemblies covering 98% of the reference genome using long-read sequences generated with Oxford Nanopore Technologies (ONT). Annotation with the sequenced transcriptome coupled with the available assembly revealed that gene predictions were more accurate, complete and contained far fewer false positives than *de novo* transcriptome assembly approaches. We sampled and sequenced the genomes and transcriptomes of 13 nematodes from Dorylaimia, Enoplia, and early-branching species in Chromadoria. These free-living species had larger genome sizes, ranging from 147-792 Mb, compared to those of the parasitic lifestyle. Nine mitogenomes were fully assembled and displaying a complete lack of synteny to other species. Phylogenomic analyses based on the new annotations revealed strong support for Enoplia as sister to the rest of Nematoda. Our result demonstrates the robustness of MDA in combination with ONT, paving the way for the study of genome diversity in the phylum Nematoda and beyond.

## Introduction

A genome reference is a prerequisite for a complete understanding of the biology and evolution of a species. Advances in long-read sequencing, together with increasing affordable costs (1), have paved the way for the ambition to study and generate genomes for the entire group of species, including every bird, vertebrate, insect, or eukaryote on Earth (2–5). However, the major challenge remains to obtain high-quality DNA and RNA from the majority of organisms across the tree of life (6). Such requirements are challenging to meet in microscopic organisms that cannot be cultured, leading to sampling bias and loss of their biological information on genetic and evolutionary studies (7–9). Recent advances in whole genome amplification (WGA) have enabled single cells or limited samples to generate sufficient DNA for sequencing (10,11), and have been applied to eukaryotic microorganisms, including fungi (9,12), marine phytoplankton (13) and parasitic nematodes (14,15) for genomic and population genetic studies (9,16).

This study investigated the feasibility of WGA combined with long-read sequencing for nematodes, which are the most abundant metazoans on Earth. More than one million nematode species are estimated to exist, but only approximately 30,000 species have been described so far (17,18). The Nematoda phylum has been classified on the basis of 18S ribosomal RNA (18S rRNA) into three lineages and five major clades: Dorylaimia (clade I), Enoplia (clade II), and Chromadoria (clade III-V). Chromadoria further includes Spirurina (clade III), Tylenchina (clade IV) and Rhabditina (clade V) as well as early derived lineages including Araeolaimida, Chromadorida, Desmodorida, Monhysterida, and Plectida (19–21). The roundworm *Caenorhabditis elegans* was the first animal to have its genome sequenced, with a size of 100.2 Mb. Since then, over 200 nematode genomes and mitogenomes of have been published (22,23). Of these, ∼72% are mainly terrestrial parasites belonging in Dorylaimia, Spirurina, and Tylenchina because of their importance in plant crop and animal health. The remaining species are terrestrial free-living nematodes in Rhabditina (17,24,25). In contrast, only one genome of a marine nematode *Plectus sambesii*, is available (26–28), despite the fact that marine nematodes comprise half of all recorded nematodes and play a crucial role in benthic communities as decomposers, predators, food sources, and bioindicators (29). Only a few marine nematode species belonging to Monhysterida and Rhabditida (30) can be cultured. Thus, obtaining enough genomic DNA for sequencing is a challenging for most of these species, making them potential candidates for the WGA techniques.

The Enoplia clade and the early derived Chromadoria lineage (19), found primarily in marine habitats, currently lack genomic data and have several important implications. Of particular interest is the phylogenetic relationship of basal nematodes, which remains unresolved due to insufficient sampling and limited resolution of 18s rRNA (20). Resolving the phylogenetic relationship of nematodes can help to understand the genomic basis of nematode diversity and the processes of evolution from free-living to parasitic life style (21,31). Increased sampling of marine nematodes and phylogenomic analyses based on mitogenomes or *de novo* transcriptomes (19,22,27,32,33) have shown better resolution of Enoplia sister to the rest of the Nematoda, but not entirely resolved (19).

Here, we have developed an assembly and annotation workflow capable of generating long genome sequences and accurate gene predictions from single nematodes. We quantified the coverage biases in the amplified sequences, produced assemblies, and assessed the accuracy of the annotations using this workflow on *C. elegans* compared to those generated *de novo* without a genome available. The finalized workflow was applied to 13 free-living marine nematodes isolated from the Taiwanese coasts. Despite obtaining lower genome coverage in these nematodes, phylogenomic analyses were performed using a total of 210,777 newly annotated genes to resolve the positions of basal clades in the Nematoda phylum. Comparisons of the genomes and complete mitogenomes of these nematodes revealed remarkable variation in genome features not observed in well-studied clades and shed light on the early evolution of nematodes.

## Materials and Methods

### Single worm DNA extraction

*C. elegans* strain N2 was grown at 22°C on NGM plates with *Escherichia coli* strain OP50 and *Aphelenchoides besseyi* APVT strain was grown at 22°C on PDA plates with *Alternaria citri*. Worms were either washed with M9 buffer from NGM plate and pelleted in 15 ml centrifuge tube or washed with the same M9 buffer for three times and starved in M9 buffer included 1% Antibiotic-Antimycotic (Thermo Fisher Scientific, Massachusetts, 15240062) in 15 ml centrifuge tube for 24 hours. A total of 13 nematode species (*Epsilonema* sp., *Enoplolaimus lenunculus*, *Linhomoeus* sp., *Microlaimidae* sp., *Mesodorylaimus* sp., *Paralinhomoeus* sp., *Ptycholaimellus* sp., *Trileptium ribeirensis*, *Sabatieria punctata*, *Rhynchonemsa* sp., *Theristus* sp., *Trissonchulus latispiculum*, *Trissonchulus* sp.) were collected in Taiwan between November 2020 to May 2022 (**Supplementary Table S1**). The sampling sites encompassed various seashores throughout Taiwan, as well as the seabed at a depth of 15-18 meters surrounding Guishan Island. Individual nematode was isolated and washed with 10% bleach for 10 seconds and transferred into with 200µl PCR tube containing lysis buffer (8µl direct PCR lysis reagent (Viagen, #102-T), 1 µl 5mg/ml Proteinase K, 1 µl 200mM DTT) and incubated in 65 ℃ for 20 minutes and 95 ℃ for 5 minutes. The DNA concertation were quantified with Qubit dsDNA HS Assay Kit (Thermo Fisher Scientific, Massachusetts, 2339927) following manufacturer’s instructions.

### Whole genome amplification

Various sources of extracted genomic DNA (gDNA) were used for amplification (**Supplementary Figure S1**): (i) Whole worm: Single worm or ten adult worms were cut into pieces with 22-gauge needle in 200µl PCR tube. In one amplification instance, single whole worm was prepared with 5% DMSO was added in polymerase mix. Genomic DNA extraction and amplification was performed with Qiagen REPLI-g Kit (150023,150043,150343, Qiagen, German). (ii) Purified DNA: single worm DNA extract with lysis buffer (8µl direct PCR lysis reagent (Viagen, #102-T), 1 µl 5mg/ml Proteinase K, 1 µl 200mM DTT) and incubate in 65 ℃ for 20 minutes and 95 ℃ for 5 minutes. The samples were further purified using Ampure XP (1:1) (A63882, Beckman Coulter, US) and diluted in 10 µl elution buffer. The multiple displacement amplification step was performed with REPLI-g Kit (150023, Qiagen) following manufacturer’s instructions.

### Genomic DNA library preparation, sequencing and assembly

For Oxford Nanopore sequencing, two digestion times were initially tested on *C. elegans* using 1.5 µg amplified genomic DNA from single or ten adult worms. Template were first digested with T7 endonuclease I (M0302L, NEB, USA) as suggested in the protocol available on the ONT community website, which was initially designed based on sequencing whole-genome-amplified genomic DNA in *E. coli* (SQK-LSK109; ver. WAL_9070_v109_revN_14Aug2019). Namely 15 minutes (as recommended in the original ONT protocol) and 30 minutes were tested. For the other nematodes, 3 µg amplified templates were digested with T7 endonuclease I for 30 minutes and subjected to library preparation according to the manufacturer’s instructions. Oxford Nanopore libraries were prepared according to SQK-LSK109 and SQK-LSK110 protocols, and sequenced on a GridION instrument. Basecalling of Nanopore raw signals was performed using Guppy (ver. 6.1.2) into a total 224.5 Gb of raw sequences at least 1 kb or longer. A summary of the sequencing data is shown in **Supplementary Table S2**. For short read sequencing, the amplified genomic DNA was sent to Biotools Co., Ltd (New Taipei City, Taiwan) for library preparation and sequencing of Illumina 150bp paired-end reads on a Novaseq 6000 sequencer.

The Flye (ver. 2.9.1) assembler (34) was used to assemble the raw ONT reads, which were then polished by four iterations of Racon (35) (ver. 1.4.11), followed by Medaka (-m r941_min_sup_g507 or r103_sup_g507, ver. 1.2.0; https://github.com/nanoporetech/medaka). The consensus sequences were further corrected with Illumina reads using NextPolish (36)(ver. 1.4.0) and haplotigs were removed using Purge Dups (37) (v1.2.5). Contigs with non-nematode origins were excluded. Genome completeness was assessed using nematode dataset of BUSCO (38) (ver. 5.1.2). Raw Illumina reads were assembled using the Spades assembler (ver. v3.14.1; spades_sc)(39). The mitochondrial genome was assembled separately using by aligning Oxford Nanopore reads to mitochondrial protein-coding gene of 52 nematode species listed in **Supplementary Table S3** using DIAMOND (40), following the approach described in (41). The circlised assembly were further annotated and manually curated using two versions of Mitos(v1.0.5) and Mitos2(2.1.0) (42).

### Single worm RNA transcriptome sequencing and assembly

The Smart-seq 2 protocol (43) was used to extract and amplify RNA from single adult worms. The resulting cDNA was sent to Biotools Co., Ltd (New Taipei City, Taiwan) for library preparation using the NEBNext® DNA Library Prep Kit (NEB, USA,20015828,20015829), and sequenced for 150bp paired-ends on an Illumina Hiseq 2500 sequencer. Individual sample statistics are provided in **Supplementary Table S2**. Sequencing reads were quality and adaptor trimmed using Trimmomatic (ver. 0.39) (44), and reads mapped to nematode ribosomal RNA genes in the NCBI database were removed. On average, 30% of the reads in each sample were identified as ribosomal RNA sequences, and after removal, 21-86% of the reads remained. *De novo* transcriptome assemblies were generated using the Spades assembler (ver. v3.14.1; option: -k=55,77)(39). The best protein encoding predictions from *de novo* assembled transcripts were produced using Transdecoder (v 5.5.0)(45) integrating homology information from the UniProt database (Release version 2021_03).

### Genome annotation

Repetitive elements were identified using RepeatModeler (ver 2.1) (46), TransposonPSI (ver 1.0.0; https://github.com/NBISweden/TransposonPSI) and USEARCH (ver 11.0)(47) based on the protocol by Berriman *et al*. (48). Repetitive DNA sequences were identified and masked using Repeatmasker (ver 4.1.2)(49). Proportions of repeat content along the non-overlapped 100kb window were calculated using BEDTools (ver 2.26.0)(50).

The proteomes of 11 representative nematodes were obtained from WormBase WBP16 (51) and are listed in **Supplementary Table 4**. Single worm transcriptome reads were mapped to the corresponding genome assemblies using STAR (ver. 2.7.7a) (52) and assembled using Trinity(ver 2.13.2; guided approach) (53), Stringtie (ver 2.1.7) (54) and Cufflinks (ver 2.2.1)(55). Transcripts generated by Trinity were mapped to the corresponding genome assemblies using Minimap2 (ver 2.1, options: - ax splice) (56), and splice junctions were quantified using Portcullis(ver 1.2.3)(56). The gene predictor Augustus (ver 3.4.0) and gmhmm(57) were trained using BRAKER2(ver. 2.1.6) (52,58) and SNAP (59) with proteomes and RNAseq mappings as evidence hints to generate an initial set of annotations. The assembled transcripts selected by MIKADO(ver 2.3.3) (60), proteome homology, and BRAKER2 annotations were combined as evidence hints for input into the MAKER2 annotation pipeline(61) to produce a final annotation for each species. Comparison of all annotation results to the reference protein-coding gene were conducted using Gffcompare (ver. v0.11.2) (62).

### Decontamination

To identify contigs of non-nematode origin, we used a combination of three methods, given the lack of uncontaminated nematode sequences in the database. First, we employed Kraken2 (ver. 2.1.2) (62) to determine the kingdom and phylum of scaffolds based on k-mers. We rebuilt the Kraken2 database to include six kingdoms and phyla: Archaea, Bacteria, Nematoda, Eukaryota (Annelida, Arthropoda, Cnidaria, Chordata, Porifera, Placozoa, Platyhelminthes), Outgroup (Human), Viruses, and Undefined. A list of species in the reconstructed database is provided in **Supplementary Table S5**. Second, we annotated genes using Braker2 and aligned them against the NCBI nr database using BLAST to assign phylum categories, such as Nematoda, Bacteria, Eukaryota, Eukaryotea-undef, Candidatus, Fungi, Planta, Viruses, Algae, Archaea, and Unclassified. Third, we aligned RNAseq reads for each species to the corresponding genome assemblies using STAR and calculated the RNAseq mapping rate of each scaffold using BedTools. We excluded scaffolds assigned to bacteria by Kraken2 and those with genes that contained 90% or more bacterial proteins. For scaffolds that could not be identified by Kraken2 or the nr database, we removed those with RNAseq mapping rates below 1,000 reads.

### Phylogenomics of nematodes

Protein data sets from 13 representative nematodes were download from WormBase WBP16 (51) (**Supplementary Table S6**). We also downloaded the assembled transcripts of one Araeolaimida, one Plectia, six Enoplia, and an outgroup Nematodmorpha from Smythe *et al* (19) (**Supplementary Table S6**). Orthogroups (OGs) were identified using OrthoFinder (63,64). Sequences in each OGs were aligned using MAFFT(v7.515)(65). A gene tree was inferred from each OG alignment using FastTree(ver 2.1.11)(66), and a species tree was inferred from all OGs gene trees using ASTRAL-Pro (67). Two sets of data were used to construct the nematode species phylogeny: (i) proteomes of nematode species downloaded from wormbase, 13 free-living nematodes sequenced in this study, and outgroup *Priapulus cauatus,* comprising a total of 27 species with 58,531 OGs (**Supplementary Table S6**), and (ii) the 27 species from (i), *de novo* transcriptomes from Smythe *et al.*, *Drosophila melanogaster, Priapulus cauatus,* and *Hypsibius dujardini*, comprising a total of 39 species with 77,411 OGs.

## Results

### Whole genome amplification facilitates sufficient DNA for long read sequencing from single nematodes

To study the genome diversity of free-living nematodes, we isolated nematodes from a variety of marine environments in Taiwan and extracted the genomic DNA (gDNA) from individual adults across 11 taxa, including the Enoplia clade, for which genome sequences are currently unavailable (**Supplementary Table S1**). Highlighting the challenge of obtaining sufficient gDNA for long-read sequencing across the Nematoda phylum, we found that yields ranged from 1.5 to 3.8 ng for individual adults and were not associated with worm size (tau = 0.2, *P* = 0.33, Kendall’s tau test, **Supplementary Table S7**). To mitigate this problem, we used multiple displacement amplification (MDA) method to amplify whole genomes from individual nematodes, yielding 6.2-51.5 µg, corresponding to approximately 4,700X and 22,000X amplification using REPLI-g mini kit, REPLI-g midi or sc kit, respectively (**Supplementary Table S2**). To assess potential amplification bias, we first sequenced the genome of the model nematode *Caenorhabditis elegans* N2. A total of 35 µg genomic DNA was obtained after MDA from an initial 1.56 ng, and sequencing using an ONT 9.4.1 flow cell yielded 7.36 Gb with a N50 of 7.74 kb, corresponding to 73.6X depth of coverage (68) (**Supplementary Table S2**).

### Whole genome amplification disparity in repetitive regions

MDA suffers from several challenges, the most important of which is highly uneven amplification (69,70). This issue can lead to incomplete genome assembly and reduced coverage in certain genome regions, and affect analyses such as copy number variants (70,71). We aligned the amplified and published unamplified ONT reads (72) against the *C. elegans* genome, and the former clearly displayed an uneven depth of coverage with 19.2% of the non-overlapping 100kb window showing less than half of the genome-wide median (**Figure 1A**). Sequencing of the amplified DNA using Illumina short reads also exhibited similar patterns, suggesting that the MDA rather than the sequencing platforms were causing the bias (**Figure S2A**). This unevenness was clearly associated with the arm-center-arm domain structures exhibited by the nematode autosomes. In the six *C. elegans* chromosomes, the arm region contains a significantly higher number of repeat sequences (*P*<0.001, Wilcoxon rank sum test) and a lower read coverage (*P* < 0.001, Wilcoxon rank sum test; **Supplementary Figure S2B,2C**) compared to the center region. However, in the X chromosome, the read coverage is more balanced due to the lower percentage of repeats in the arm region (Median of repeats 16-21% vs. 15-39% in autosomes; **Supplementary Figure S2D**).

**Figure 1.**
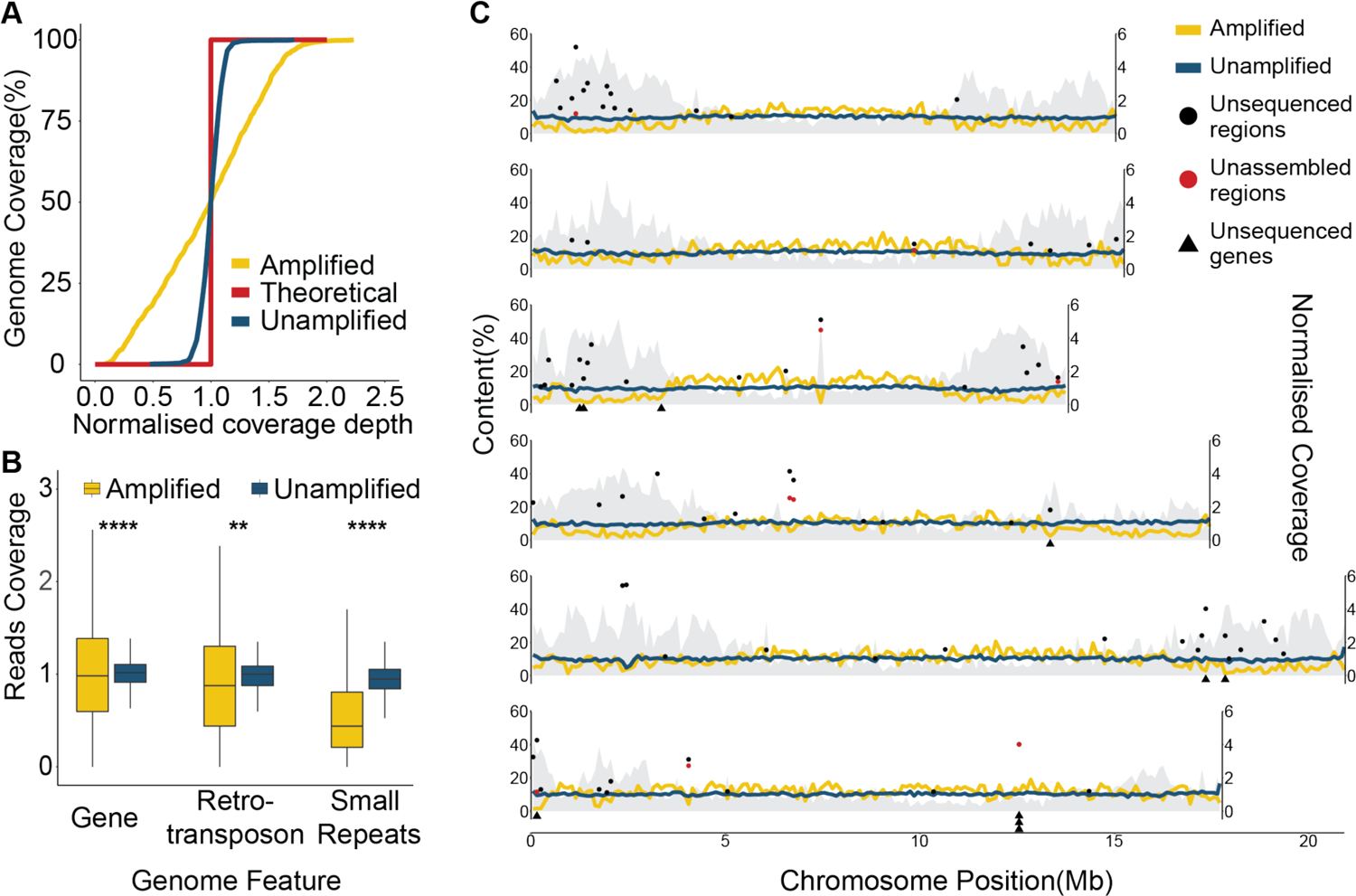
Sequencing coverage of *C. elegans* genomic DNA. **A.** Cumulative genome coverage versus genome wide median. The red line indicates the theoretical coverage of unbiased coverage. More derivation away from this line suggest less uniformity across the genome. **B.** Normalized read coverage on genes and repeats. Repeats were categorised into two groups: retrotransposons and small repeats. Retrotransposons include long interspersed nuclear elements (LINEs), short interspersed nuclear elements (SINEs), and long terminal repeats (LTRs). DNA transposons, RC Helitron, rRNA, snRNA, tRNA, satellite, simple repeat, and unknown repeats were labeled as small repeats. **: *P* < 0.01, ****: *P* < 0.0001. **C.** Lines represent normalized read coverage of amplified and unamplified data. The black and red dots represent the top 10% regions (11.1-44.8kbp and 10.1-54.5 kbp) that were not sequenced and not assembled. The triangles represent the position of genes that were not sequenced at all. The shaded area indicates the proportions of repeats along the chromosomes.

We calculated the proportions of 17 genomic features in 100kb non-overlapping windows and found that the presence of rolling-circle transposable elements (RC/Helitron) contributed least to sequence coverage (R^2^=0.36,*P*< 22e-16, Pearson test), followed by unclassified repeats, DNA transposable elements and Satellite (**Supplementary Figure S3**), suggesting that the reduced coverage displayed at autosome arms were mainly due to enriched repeats (68). Within repeat class, small repeats were the primary repeat class affecting coverage (**Figure 1B; Supplementary Figure S4**). Of the 6,332 rolling-circle transposable elements regions totaling 1,409,947 bp, on average a reduced 36.7% of coverage compared to the genome median (68X vs. 25X; **Supplementary Figure S4**). The coverage of amplified data on the gene, retrotransposon, and other small repeats were significantly lower than unamplified data (**Figure 1B**). Finally, we determined 0.49 Mb that were not sequenced at all ranging from 2 to 40,137 bp. As expectedly, 17% were repeats and 80 (0.3%) of genes were affected. Ten genes located at near the arm of III, IV, V, and X chromosome were not completely sequenced (**Figure 1C**). At the exon level, 0.09% were have also been affected, including 96 not sequenced CDS.

Due to differences in repeat content between species, we sought to evaluate the impact of MDA by analysing the amplified reads from the genome of the plant parasitic nematode *Aphelenchoides besseyi*, which has a smaller genome (44.7 Mb) and lower repeat content (5.38%) compared to other nematodes (73). A similar but less pronounced pattern of uneven coverage was observed in this nematode compared to *C. elegans*. (**Supplementary Figure S5A**). In the *A. besseyi* chromosomes, the chromosome arm also contains significantly higher percentage of repeats sequence and lower reads coverage (*P* < 0.001, Wilcox test; **Supplementary Figure S5B, 5C**). While we observed reduced coverage in repeat regions, we were able to capture the entire genome with only 0.06% (29,230 bp) of the genome not sequenced, with missing regions ranging from 1 to 2,540 bp and affecting only six repeats and four genes. Taken together, these observations suggest that the MDA approach is a robust approach for capturing the entire nematode genome, although additional sequencing may be required to rescue repetitive regions in some species.

### Longer T7 endonuclease digestion time increase ONT sequencing performance

To enhance the yield and minimize the bias of sequencing amplified samples, we attempted to optimise different stages of the workflow (**Supplementary Figure S1**). One of the challenges was that the branching templates generated by MDA were unsuitable for Oxford Nanopore sequencing as they can block sequencing pores and reduce sequencing yield (74). T7 endonuclease I was used in the existing protocol to generate linear templates by cutting the junction of the branching template. We found that the digestion time of T7 endonuclease I affected the dropout rate of sequencing pores and sequencing output (**Supplementary Figure 6A**) (75). A longer digestion time (30 versus 15 mins) improved the sequencing performance by reducing the dropping rate of sequencing pores and increasing the sequencing yield (**Supplementary Figure 6B, 6C**). In addition, we tried three approaches to address the influence of repeats that reduce amplification efficiency (**Supplementary Figure S1**), but no improvement was observed (**Supplementary Figure S2A**), indicating that uneven coverage is more related to the polymerase efficiency in amplifying repeats.

### Complete genome assemblies from amplified sequences

To evaluate the feasibility of generating assemblies from amplified sequences, we generated genome assemblies based on different data types and sources (**Supplementary Table S8**). On average 112.5X Illumina and 32.6X-88.6X ONT reads were used and the initial assemblies produced under the default options yielded were 72.5-115.9Mb and the recently updated size of 100.2Mb (76), suggesting that the biased genome coverage in the amplified reads remains a challenge in the assembly process. The final assemblies were produced using the meta option of the Flye assembler, with haplotigs removed, screened for contamination and polished using Illumina reads (**Table 1;** Methods), resulting in more similar genome sizes (100.9 to 98.3 Mb, **Table 1**). Compared to the assembly from unamplified long sequences (72), the N50 of the genome-amplified assembly is 20% shorter than the unamplified data, presumably due to the shorter ONT sequence length achieved by MDA (**Table 1**).

**Table 1.** Statistics of *C. elegans* genome assemblies

|  | Reference | Amplified | Unamplified |
| --- | --- | --- | --- |
| **Reads N50(Kb)** | - | 7.7 | 21.1 |
| **Depth** | - | 73.5 | 88.6 |
| **Size (Mb)** | 100.2 | 100.9 | 103.8 |
| **Seq number** | 7 | 484 | 79 |
| **Longest (Mb)** | 20.9 | 2.7 | 15.1 |
| **Minimum(bp)** | 13,794 | 1,041 | 815 |
| **N50(Kb)** | 17,494 | 614 | 6,740 |
| **L50** | 3 | 45 | 5 |
| **N90(Kb)** | 13,784 | 126 | 2,691 |
| **L90** | 6 | 197 | 15 |
| **BUSCO completeness (%)** | 98.7 | 96.8 | 92.8 |
| **Assembly covered (%)** | - | 98.0 | 99.7 |

We assessed the completeness of the *de novo* assemblies of single and ten pooled nematode(s) by first aligning the contigs back to the reference. The unamplified data covered 99.7% of the reference genome, compared to 98% for the single-worm amplified assembly (**Table 1**). 2.5 - 4.4 Mb of the reference genome was not covered by the amplified genomes. We benchmarked the completeness of the assemblies using universal single-copy orthologs (BUSCO,(38), which were similar regardless of amplification (**Table 1**). The unassembled regions coincided with not sequenced or highly repetitive regions which were mostly located on the chromosome arms (**Figure 1C**). Taken together, the results show that the capability of sequencing the genome of the nematode with only a single worm using the WGA method is equivalent to using multiple worms. Interestingly, we observed a decrease in reference coverage (98.0 vs. 95.8%) and BUSCO completeness (96.8 vs. 95.1%) as the number of worms increased from single to ten worms prior to MDA. In addition, more contaminated sequences were present (17.1 Mb vs. 0.6 Mb in single worm).

### High quality annotations from single nematode genome and transcriptome

To quantify the difference between annotation based on a single worm genome and transcriptome, we profiled the transcriptome from a single *C.elegans* adult based on the Smartseq2 protocol and generated ∼10Gb of Illumina reads (43); **Supplementary Table S2**) We generated annotations either by *de novo* assembly of these reads or mapped these reads to genome assembly and used these reads as evidences in the MAKER pipeline (61). To evaluate the accuracy of the annotations from different approaches and datasets, all gene predictions were aligned back to the *C. elegans* reference and compared to the most recent annotation of 20,184 protein encoding genes from WormBase (ver WBP14, (51); **Figure 2** **and Supplementary Table S9**). Comparing two versions of the *C.elegans* reference (WBP14 vs.WS100), more loci (0.6% vs 9.1%) were annotated in the current reference, demonstrating the improventment in gene annotation over time. Gene structure predicted using single worm RNA-seq mapped to the reference genome by Stringtie (54) had lower sensitivity and precision compared to the reference proteome, and the MAKER pipeline produced a more accurate prediction when using these mappings as hints. More importantly, the same pipeline produced predictions with only a slightly reduced accuracy of 1.2% on average when the genome-amplified assembly was used instead (**Figure 2A**). As expected, these annotations were more sensitive and accurate than ones that was originally published (WS100) and *de novo* transcriptomes for all metrics (**Supplementary Table. S9**). The *de novo* transcriptome had more missing exons (36.6% vs. 7.0%-23.9%), missing loci (35.7% vs 2.5-23.9 %) and fewer matching transcripts (7,227 vs 10,163 - 13,294)(**Figure 2B, 2C**). Of particular note, the proportions of 50% and 95% assembled genes, which are imparative for phylogenomic analyses, were 76.8% and 48.3% lower in the *de novo* transcriptome compared to the annotation produced with the available genome-amplified assembly (**Supplementary Table. S9**). These results indicate that the genome-amplified assemblies can be annotated with reasonable accuracy using the existing pipeline.

**Figure 2.**
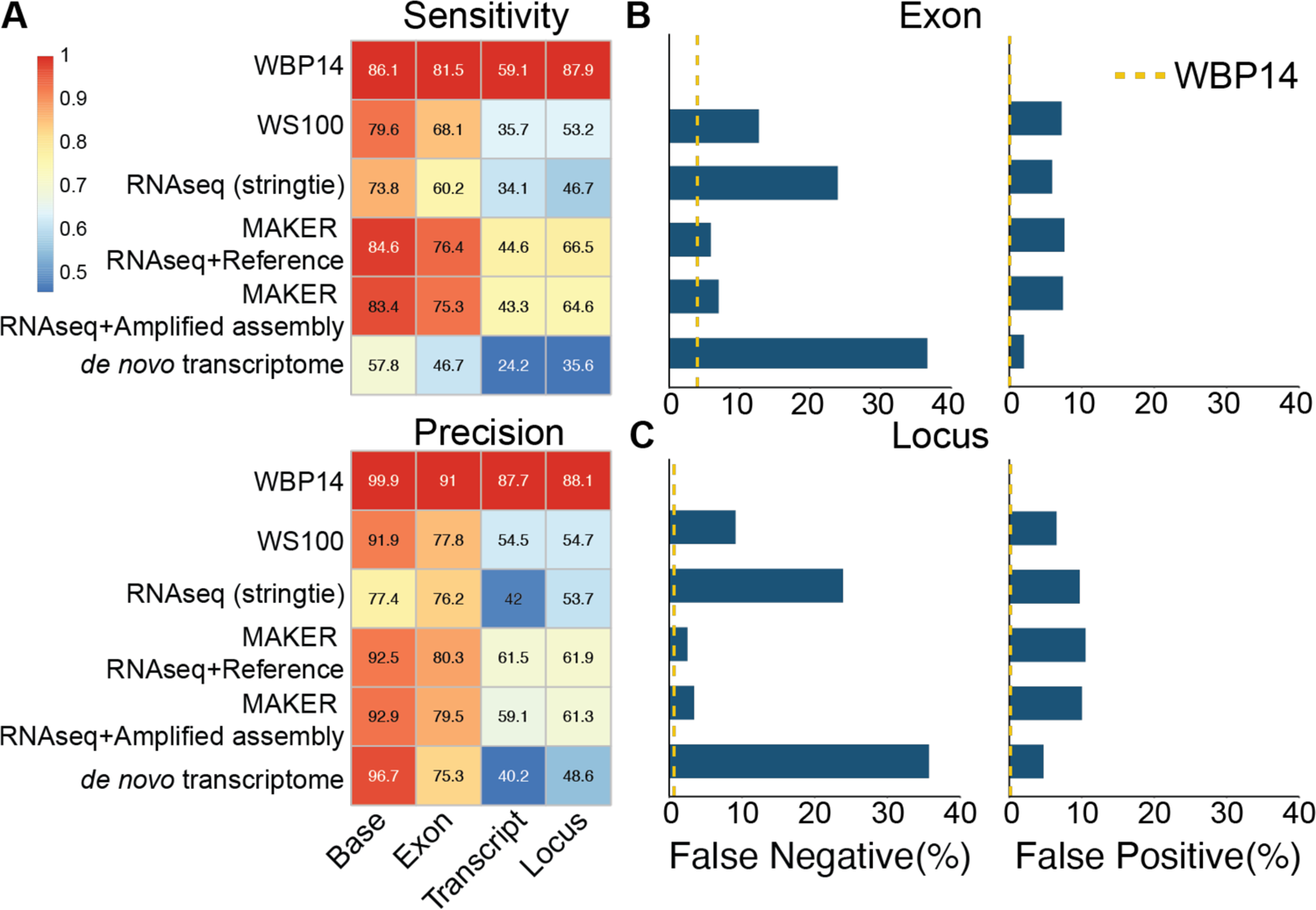
Comparison of annotations using different approaches and datasets. **A.** Sensitivity and precision on base, exon, transcript and locus level. WBP14 indicates the baseline performance when the reference proteome was aligned back to the reference genome using Minimap2. Colour of the heatmap is the value of each sample divided by the values in the WBP14 comparison. **B,C.** Percentage of false negatives and false positives in exon and locus. False negatives is the precentage of reference genes missing from the predictions, whereas false positives are new genes in the predictions that are not present in the reference proteome. Yellow dash line represent the value of the WBP14 comparison as baseline.

### Genome characteristics of free-living nemtodes

We applied our optimised sequencing and annotation protocol to 13 free-living nematodes from three clades collected from the north coast of Taiwan (**Table 2**). On average, 12 Gb of ONT genomic reads and 5 Gb of transcriptome reads were sequenced from two adults per species (**Supplementary Table S2**). Strikingly, the assemblies ranged from 147.9 to 792.4 Mb indicating that the genome size of free-living nematodes is larger than that of most published parasite genomes. In particular, nematodes belonging to Enoplia tend to be larger than other clades (**Figure 3A**), with the 792.4Mb assembly of *T. latispiculum* being the largest currently recorded in nematodes. Some of the assemblies may underestimate the true genome size, as the sequence coverage was 11-80X, warranting additional sequencing. Using the MAKER pipeline, 22,422 - 59,888 protein-coding genes were annotated in 13 free-living nematode genomes and were 34.5 to 92.6% complete based on BUSCO analysis (**Supplementary Table S10**). The intron distribution of the Dorylaimia and Chromadoria lineages sequenced in this study had a similar pattern, peaking around 50 bp (**Table 2, Supplementary Figure S7**) and were similar to previously published nematodes in these clades (77). Interestingly, there are fewer but longer introns in the four Enoplia species, suggesting a different intron distribution in the last common ancestor of this clade compared to the rest of nematodes (**Figure 3B, Supplementary Figure S8**). Orthology inference using Orthofinder (20, 21) placed these gene models and those of 13 other nematode genomes into 54,890 orthologous groups. Within these orthologous groups, 45.9 % (25,218 orthologous groups) were shared between two or more species, consistent with previous observations of extensive clade-specific families in nematode lineages (78). In addition, 16-69% of the genes in 13 free-living nematode species in were species-specific.

**Figure 3.**
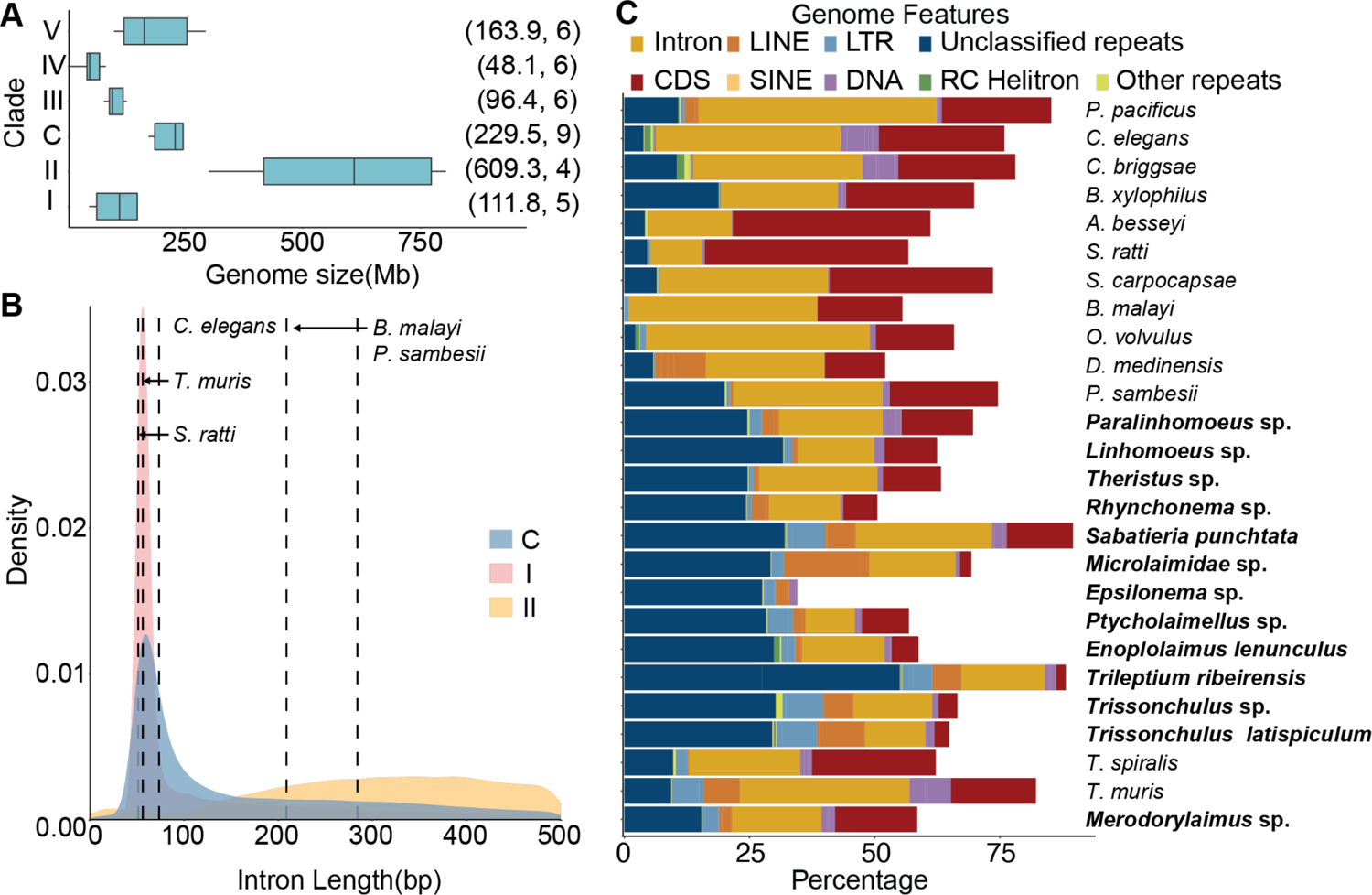
Nematode genome size, intron distribution and genome structure. **A.** Genome size variation between different nematode clades. The brackets denote the median genome size and number of nematodes used for analysis, respectively. **B**. Intron distribution of nematodes. Dashed lines are the intron median of the representative species in clades I, III, IV, V and Chromadoria(C). **C.** The proportion of different genome features in the nematode genomes. Bold letters represent the nematode genome assembled in this study. The ‘Other repeats’ feature include the sum of tRNA, snRNA, rRNA, simple repeats and satellite.

**Table 2.** Genome statistics of 13 free-living nematode genomes assembly.

| Species | clade | Size (Mb) | Seq (num.) | Longest (Kb) | N50 (Kb) | Gene (num.) | Intron length median(bp) |
| --- | --- | --- | --- | --- | --- | --- | --- |
| *Mesodorylaimus* sp. | I | 148 | 1,727 | 2,531.2 | 409.2 | 16,194 | 64 |
| *Trissonchulus* sp. | II | 447 | 21,105 | 252.3 | 36.3 | 14,926 | 538 |
| *T. ribeirensis* | II | 746 | 30,783 | 291.3 | 42 | 12,305 | 1,796 |
| *E. lenunculus* | II | 300.9 | 4,131 | 891 | 164.7 | 10,199 | 564 |
| *T. latispiculum* | II | 792.4 | 32,069 | 326.8 | 40.5 | 20,486 | 437 |
| *Theristus* sp. | C | 170.7 | 5,900 | 427.2 | 58.1 | 15,115 | 248 |
| *S. punctata* | C | 390.8 | 8,097 | 959.4 | 90.9 | 23,376 | 98 |
| *Ptycholaimellus* sp. | C | 238.3 | 12,730 | 319.1 | 33.3 | 15,899 | 58 |
| *Linhomoeus* sp. | C | 240.5 | 14,458 | 316.5 | 28.9 | 20,474 | 88 |
| *Paralinhomoeus* sp. | C | 177.3 | 3,869 | 553.7 | 80 | 18,659 | 76 |
| *Microlaimidae* sp. | C | 575.8 | 35,431 | 240.1 | 24.1 | 14,273 | 981 |
| *Epsilonema* sp. | C | 171.9 | 7,781 | 250.3 | 36.9 | 17,522 | 245 |
| *Rhynchonema* sp. | C | 222 | 9,378 | 328.1 | 43.5 | 11,349 | 211 |

Of of the 13 nematode species sequenced in this study, nine were able to assemble a circular mitochondrial genome (mitogenome) with a read coverage depth of 39-2,656X. Interestingly, only five species be completely predicted all 12 proteins typically found in nematode mitogenomes. These mitogenomes are highly rearranged compared to other clades (Rhabditina, Spirurina, and Tylenchina) (**Supplementary Figure S9**). The gene order in *Mesodorylaimus* sp. showed lack of synteny compared to other species in Dorylaimia, consistent with previous observations of a high rearrangement rate in the Dorylaimia species (79).

The free-living nematodes contained significantly more repeats than nine representative parasitic nematodes across three clades (*P*<0.001, Wilcoxon ranked sum test; **Figure 3C** and **Supplementary Figure S10A**). This was particularly evident in Enoplia and Chromadoria, which contained a significantly higher proportion of repeats (24.1% to 69.5%) in their genomes compared to parasites (0.8% to 31.4%). In contrast to most published genomes in the Dorylaimia and Rhabditina clades, which were enriched in DNA transposons (78), (**Supplementary Figure S10B**), LTR or LINE repeats were more abundant in six free-living nematode species, especially in the two *Trissonchulus* species (7.4-6.8% vs. 1.2-5.4% other species), both LTR and LINEs are enriched (**Supplementary Figure S10C-E**). A total of 16,631 unknown repeat families were identified in the 13 nematodes, which were clustered into 16,557 sequence groups based on 90% identity suggesting that they were species specific. These unclassfied repeats were evenly distributed across the genome with the exception of three species in Enoplia (*Trissonchulus* sp. *T. latispiculum and T. ribeirensis*), which each had a dominant family comprising 2.2-5.6% and 0.7-1.5% of the unclassified repeats and genomes, respectively.

### Enoplia is sister to the rest of the nematode classes

To resolve uncertainties in the basal branch order of the Nematoda phylogeny, in particular the relative placement of Enoplia and Dorylaimia, a species tree was inferred based on a coalescent-based analysis (67,80) of 58,531 paralogous gene trees from 26 representative nematode species and the cactus worm *Priapulus cauatus* as an outgroup. The species phylogeny separated nematodes into six groups, including five clades and groups of the early derived Chromadoria lineage (20), and placed Enoplia as a sister group to Dorylaimia and Chromadoria lineage, both with strong boostrap support (**Supplementary Figure S11**). The topology remained similar when we included an additional nine *de novo* transcriptomes from Enoplia and Chromadoria and a further three outgroups (**Figure 4**) (19). The combined phylogeny shows that nematodes can be separated into the previously designated five clades (20) and support for the placement of Enoplia remained robust (Astro-pro, bs=100). In the early derived Chromadoria lineages, most of the lineages were grouped by order, with the exception of the Monhysterida lineage which was paraphyletic and separated by Areaolmida.

**Figure 4.**
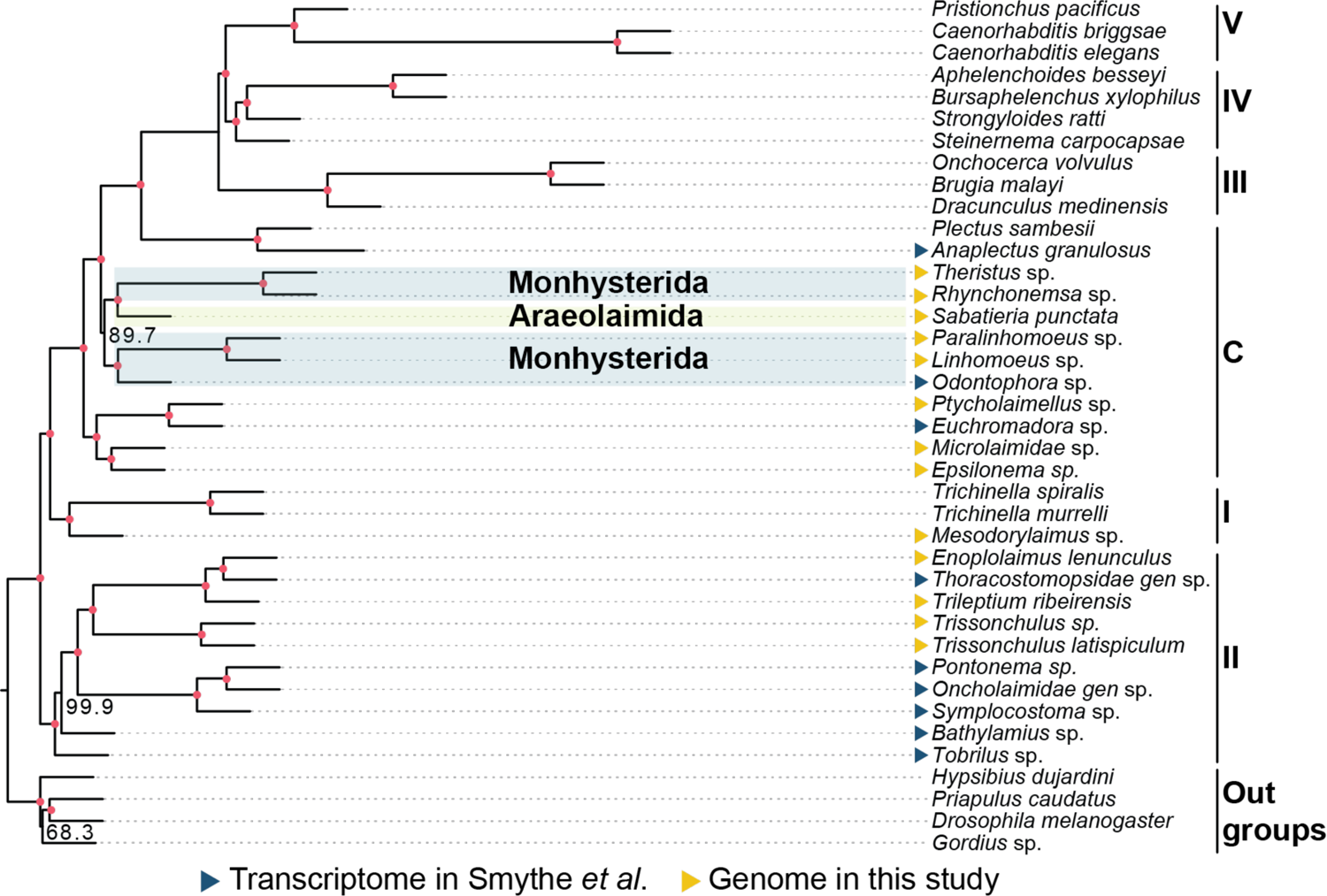
Phylogenetic analysis combining nematode transcriptome and genome. The bold font represents the Nematoda order in the shade area. The Roman numerals on right represent the five clades of Nematoda and C represent the basal Chromadoria lineage. The colours of triangles denote data source. The red dots in branches denote a bootstrap support value of 100.

## Discussion

In this study, we demonstrated the feasibility of generating genome assemblies from single adult nematodes using multiple displacement amplification. By testing the protocols on *C. elegans,* we were able to fully quantify the extent of bias and address it with existing analysis pipelines. With a genome size of 148-792.4 Mb in 13 nematodes, sequencing on a single MinION flowcell can be expected to provide approximately 37.8X depth of coverage. We demonstrate that a genome assembly and accurate gene annotations can be achieved with this workflow and further sequenced the genomes of 13 free-living nematodes. Of these genomes, four are the first reported in the Enoplia clade revealing their unusually large genome sizes and structures (**Figure 3**). Through phylogenomics, we established Enoplia as sister to the phylum Nematoda, suggesting a marine origin in the last common ancestor of nematodes (19). We overcame the stage of obtaining axenic cultures (8) whilst assembly and annotation can be achieved within two weeks of nematode isolation. Assuming that 1ug is required for long-read sequencing, combining MDA with ONT sequencing thus provides a cost- and labour-effective solution (1) to generate complete assemblies in organisms with as little as 50 picograms of starting material.

The advantage of using a single individual for whole genome sequencing is also seen in the sequencing of organisms such as obligate symbionts and helminth eggs, where it is possible to overcome obstacles such as inability to culture and inaccessibility in the live host (9,81). The use of a single nematode had several benefits over pooling multiple worms, for instance closely related nematode species have imperceptible morphological differences that increased the risk of mixing different species (82,83). For example, a host can be infected with multiple *Anisakis* species with no morphological differences (84). In addition, natural populations are likely to have high levels of heterozygosity, which also affects the quality of assembly and annotation, as observed in this study.

The MDA method used in this study is known to result in uneven read coverage (85). This unevenness is thought to be caused by the formation of secondary structures that reduce the efficiency of the phi29 polymerase used in the amplification process, particularly in repetitive sequences that are prone to forming such structures (81, 82). Despite these challenges, only a small portion of the *C. elegans* genome remained unsequenced, including ten genes representing only 0.4% of the genome. This approach allowed us to effectively assemble the genome, with only 2.5% missing due to the combined challenges posed by repetitive sequences and reduced coverage. One existing limitation was that the N50 of the amplified data was capped to ∼8kb, resulting in a more fragmented assembly compared to assembly from unamplified sample. Nevertheless, the BUSCO completeness values suggest that the amplified assembly is complete and capable of generating high-quality annotation compared to reference genome (96.8% vs 98%) (86). In addition, we show that gene prediction is superior when a complete genome is available compared to *de novo* transcriptome assemblies especially in the number of 95% assembled loci (82.9 vs 40.0%), and is ideal for subsequent phylogenomics and comparative genomics analyses.

To conclude, we demonstrate the feasibility of incorporatng whole genome amplification into investigation the study of microbial biodiversity from sampling to comparative genomic analysis. By thoroughly characterising and accounting for the inherent nature of this approach, complete assemblies and accurate gene predictions can be generated. The availability of the new free-living genomes has allowed us to address outstanding questions and offer new biological insights. As long-read sequencing advances in accuracy and affordability, we envision that a complete assembly will be available for any species that were once considered inaccessible.

## Data Availability

All sequences generated from this study were deposited on NCBI under BioProject PRJNA953805 and accession numbers of the free-living nematodes can be found in **Supplementary Table S3**.

## Authors’ Contributions

IJT conceived and the study; YiCL, YuCL, MCW and YCT carried out the sampling; YiCL isolated the nematodes; YiCL and HMK conducted the experiments and ONT sequencing; YiCL performed the assemblies and annotations with help from HHL and YuCL; YiCL carried out analyses. YiCL and IJT wrote the manuscript with input from TK and others. All authors read and approved the final manuscript.

## Acknowledgements

We thank Wei-An Liu for testing out the initial MDA protocols in yeast. We thank NGS Genomics core lab of Academia Sinica of Taiwan for sequencing the initial single worm RNAseq data. We would like to thank the National Center for High-performance Computing (NCHC) of the National Applied Research Laboratories (NARLabs) of Taiwan for providing computational resources and storage resources.

