## Supplementary Figures for "Single worm long read sequencing reveals genome diversity in free-living nematodes"

**Figure S1. Schematic diagram showing the strategies we used to optimise the whole genome amplification (WGA) step.** Four types of input material were used: single worm, pooling ten worms, single worm with 5% DMSO, and gDNA extract from single worm. After WGA, templates were sequence with Illumina sequencer. Template digest with different T7 end I treatment times were sequenced with Oxford Nanopore platform.

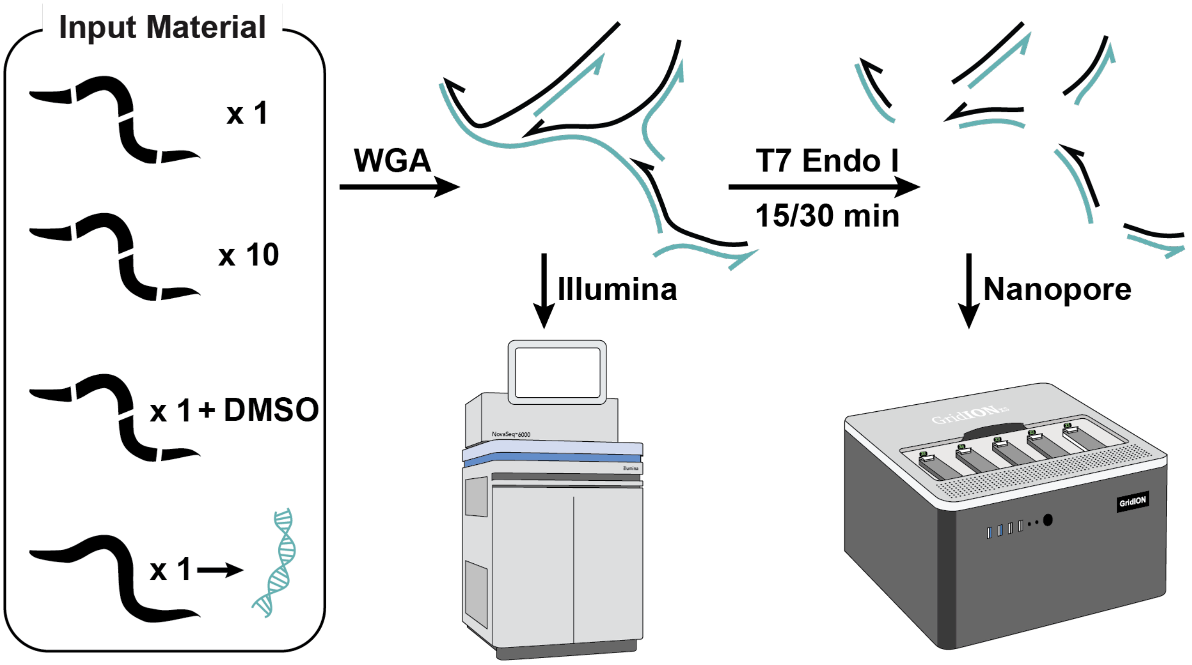

**Figure S2. Repeat content and sequencing coverage in *C. elegans***

**A.** Cumulative Illumina read genome coverage versus genome wide median in *C. elegans*. **B.** Proportion of repeat content in arm-center-arm region of *C. elegans* six chromosomes. **C.** Normalised read coverage in arm-center-arm of six chromosomes. *: $P< 0.05$ ***: $P < 0.001$, NS.:$P > 0.05$. **D.** Cumulative Nanopore read genome coverage versus genome wide median in six chromosomes of *C. elegans*.

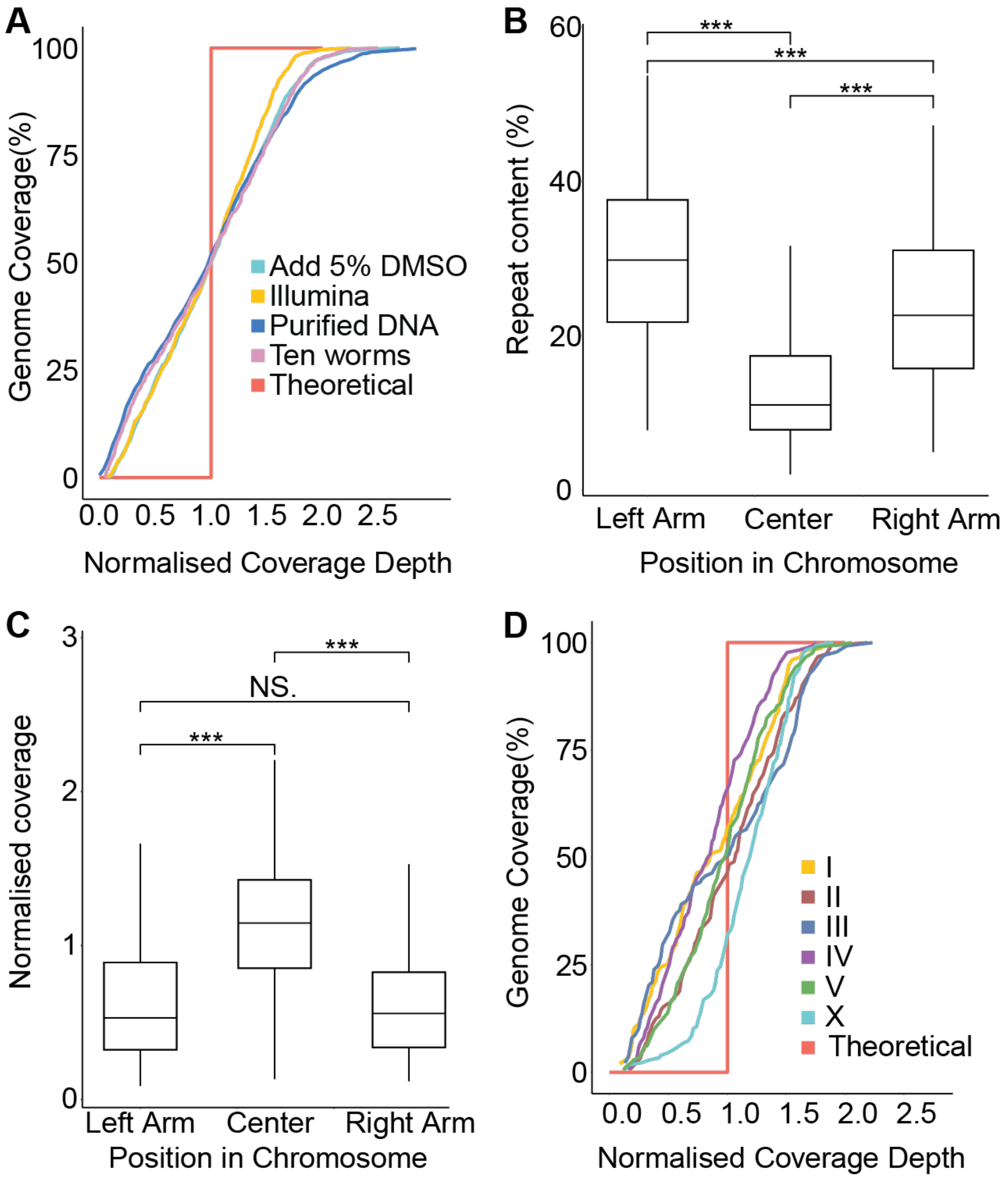

**Figure S3. Correlation between repeat content and sequence coverage in non-overlapping 100kb windows.** Scatter plots showing read coverage of different genome features, including repeats and genes in amplified samples. Genome coverage in repeats, including RC Helitron, and DNA transposons, and unknown repeats were siginificantly negative correlated.

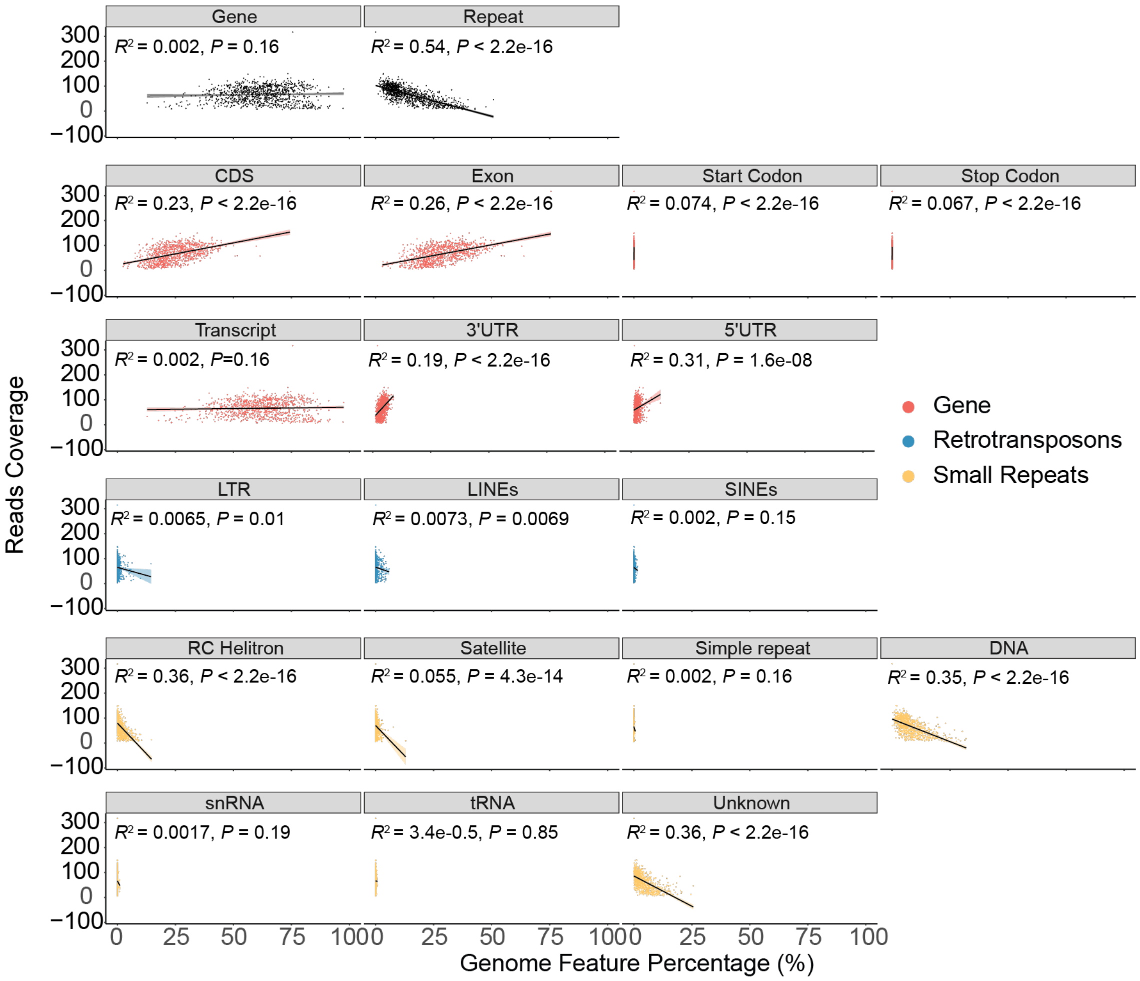

**Figure S4. Reads coverage across genome features.** The box plot shows the normalised reads coverage on each genome feature. RC Helitron had the lowest coverage. The numbers box represent the median coverage of each feature.

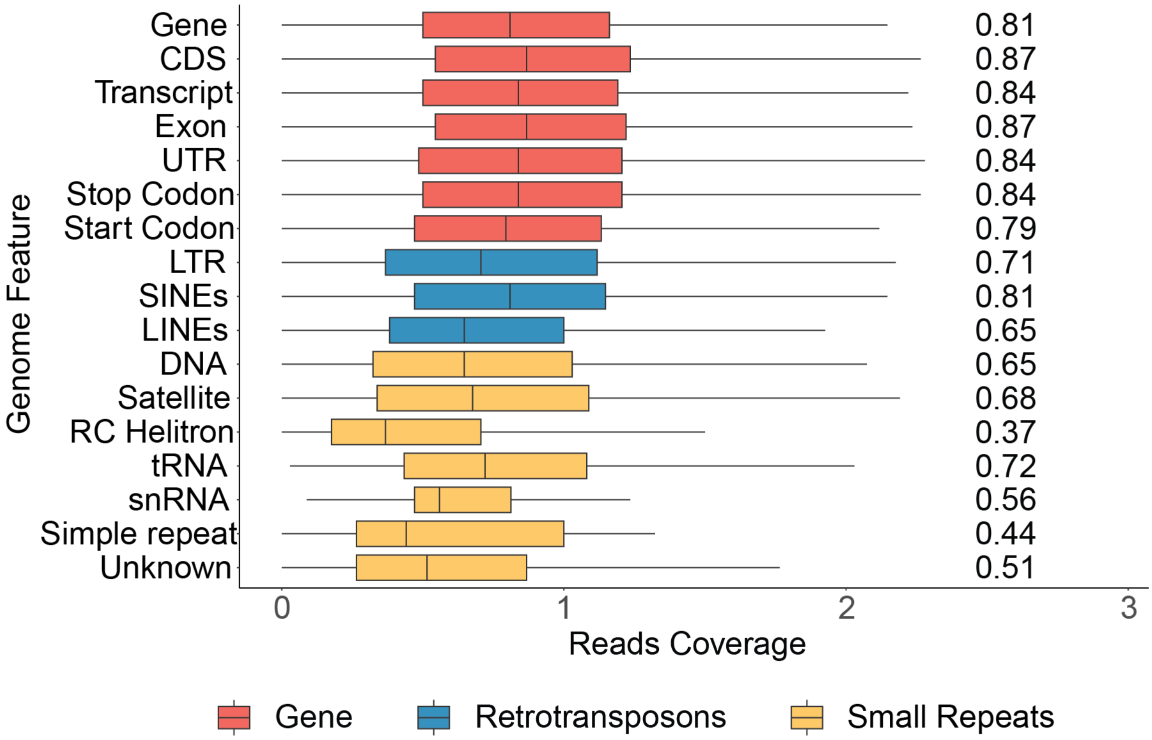

**Figure S5. Coverage uneveness of WGA on *A. bessyi*. A.** Cumulative Nanopore reads genome coverage versus genome wide median in six chromosomes of *A. besseyi* genome. **B.** The repeat cotent in *A. besseyi* chromosome arm and center. **C.** Sequencing reads coverage in *A. besseyi* chromosome arm and center. *: $P < 0.05$ ***: $P < 0.001$, NS.:$P > 0.05$.

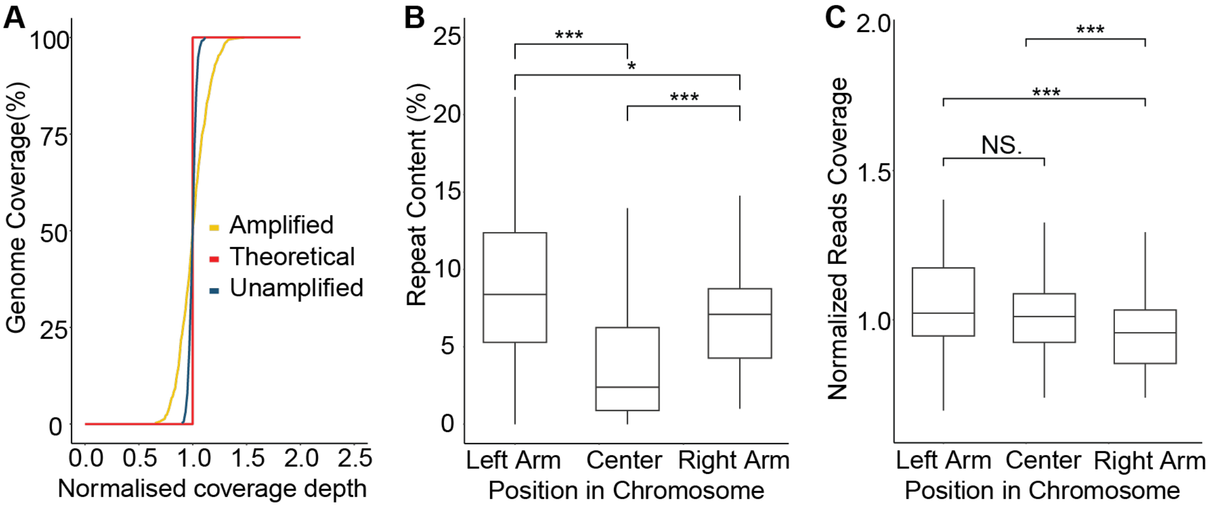

**Figure S6. Effect of T7 digest times on the Oxford Nanopore sequencing platform. A.** Number of sequencing pores over sequencing time. The triangles represent the time of mux scan. **B.** Read length distribution from a subset of 6.6Gb data from each sample. **C.** Cumulative sequencing output (Gb) over time. Reads length distribution showed that with 15 minutes digestion, the reads length is generally shorter. This showing longer digestion time indeed generate more linear template and have less defect on sequencing.

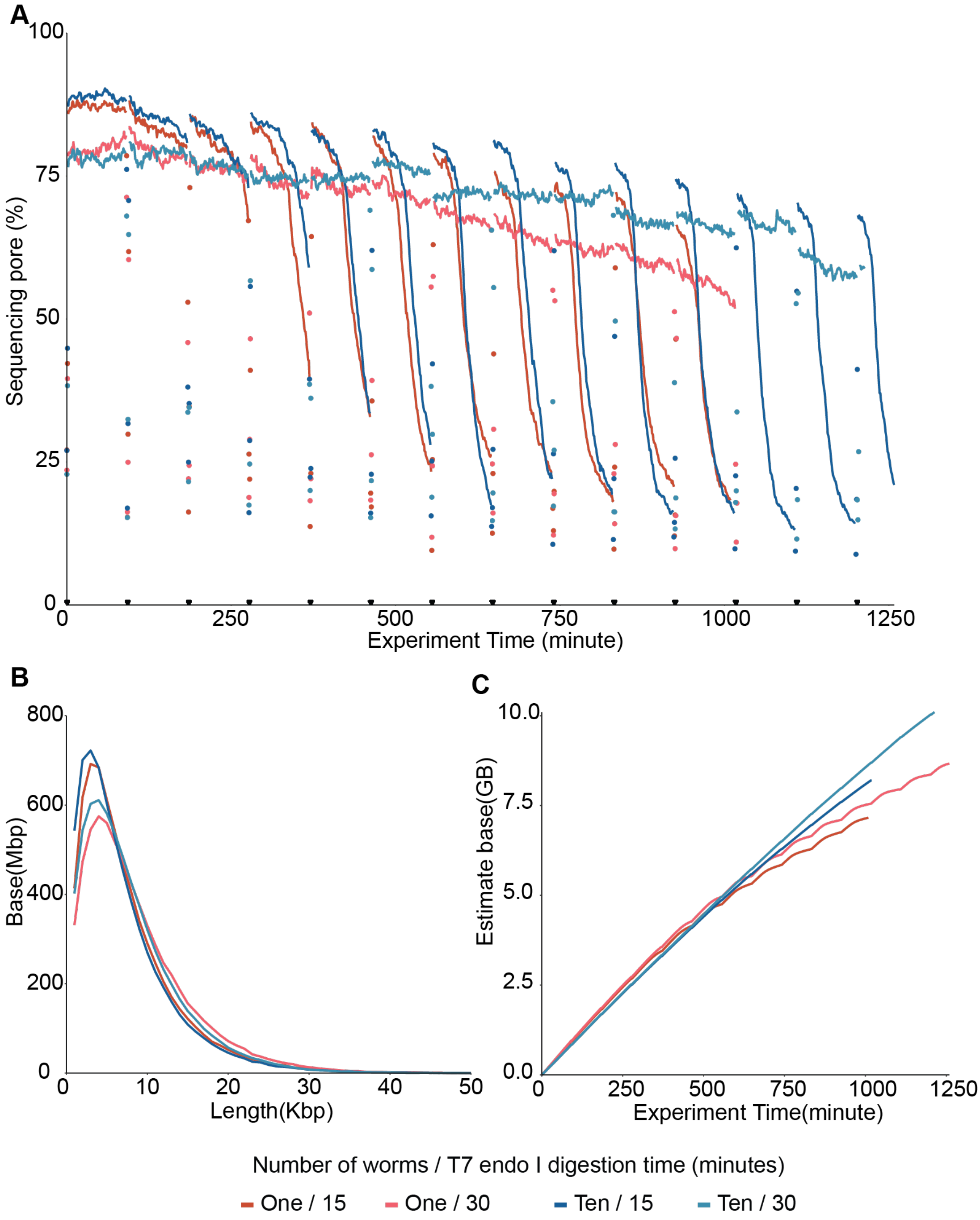

**Figure S7. Distribution of intron length in nematodes.** Different colours denote different Nematoda clades.

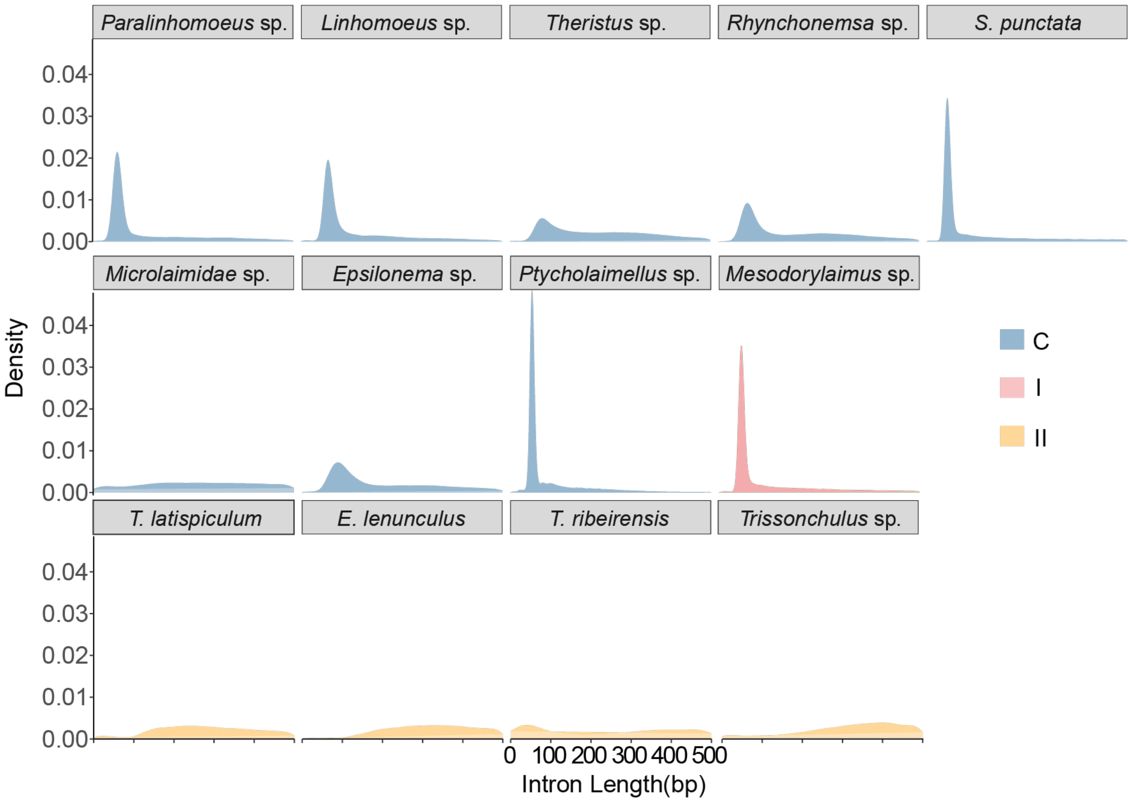

**Figure S8. Distribution of intron length (A) and numbers (B) in nematode proteomes.** Different color denote different Nematoda clades.

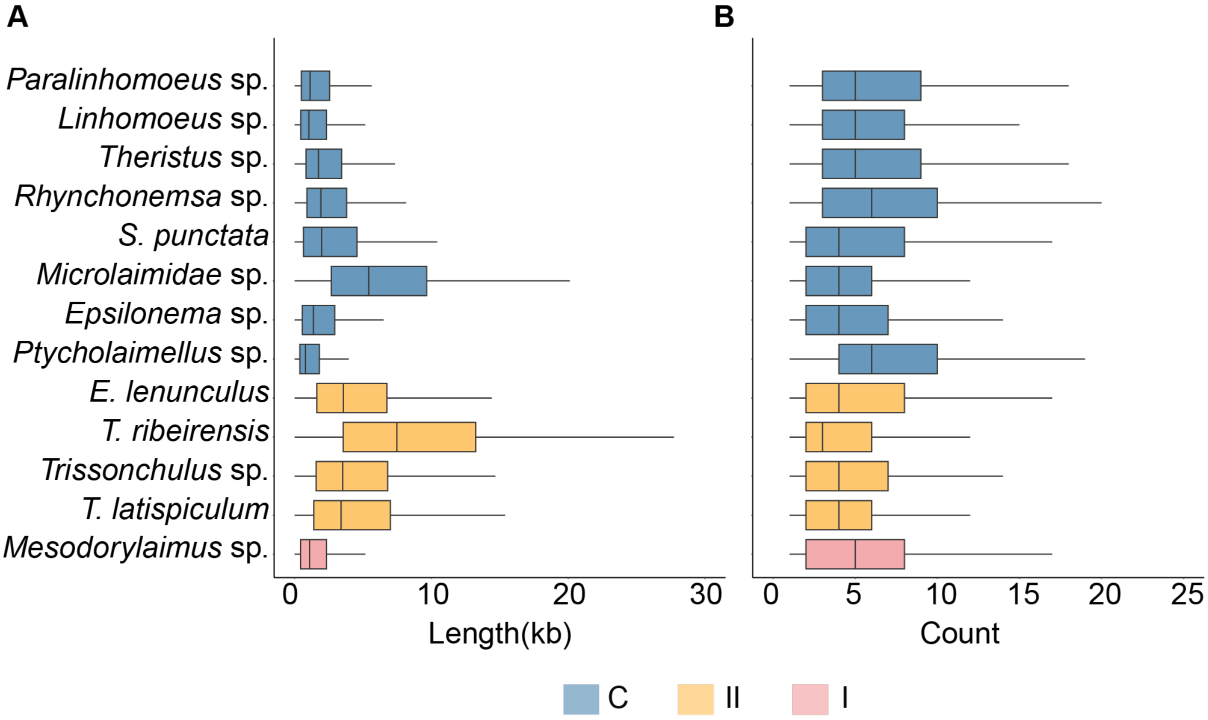


**Figure S9. Mitochondrial genes order in nematodes.** The mitochondral gene order from representative published nematode mitogenomes in clades C and I, and nine species with near complelete assembled mitogenome in this study, are denoted by bald letters.

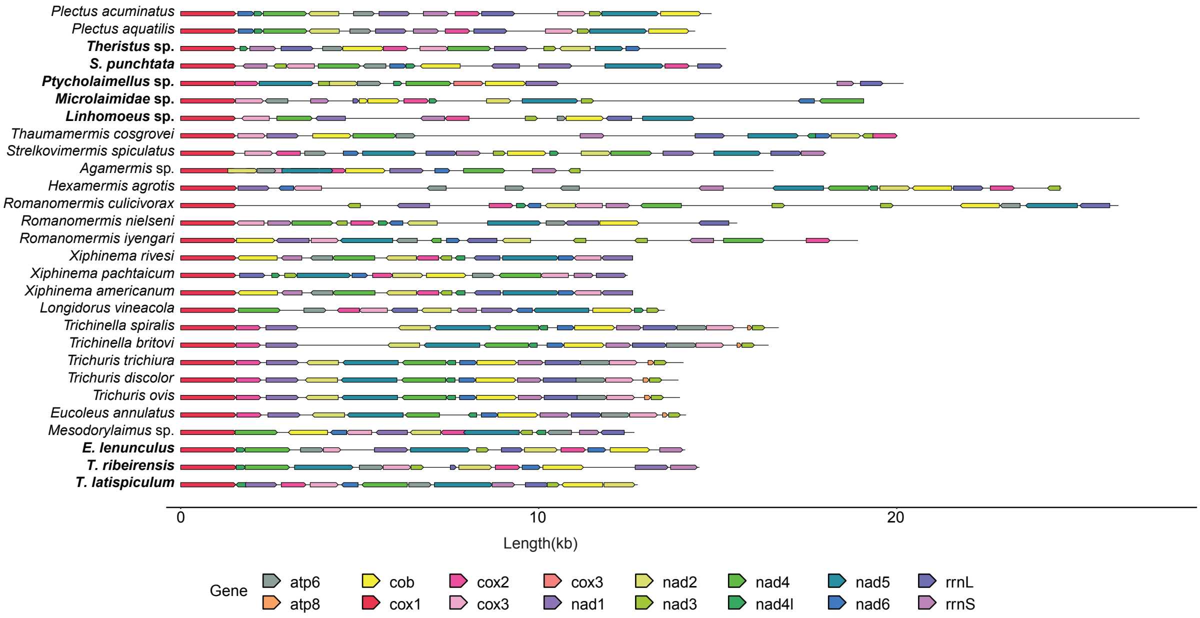

**Figure S10. Repeat contents in nematode genomes. A**. Proportions of repeats in free-living and parasitic nematode genomes. **B.** The proportions of different repeat feature, coloured by percentages. **C-E.** Proportions of LTR, DNA, LINEs. Free-living nematode had significanly more repeats than parasitic nematode. The pattern of enrichment of DNA transposon, LTR, and LINE families appeared to be specific to each nematode.

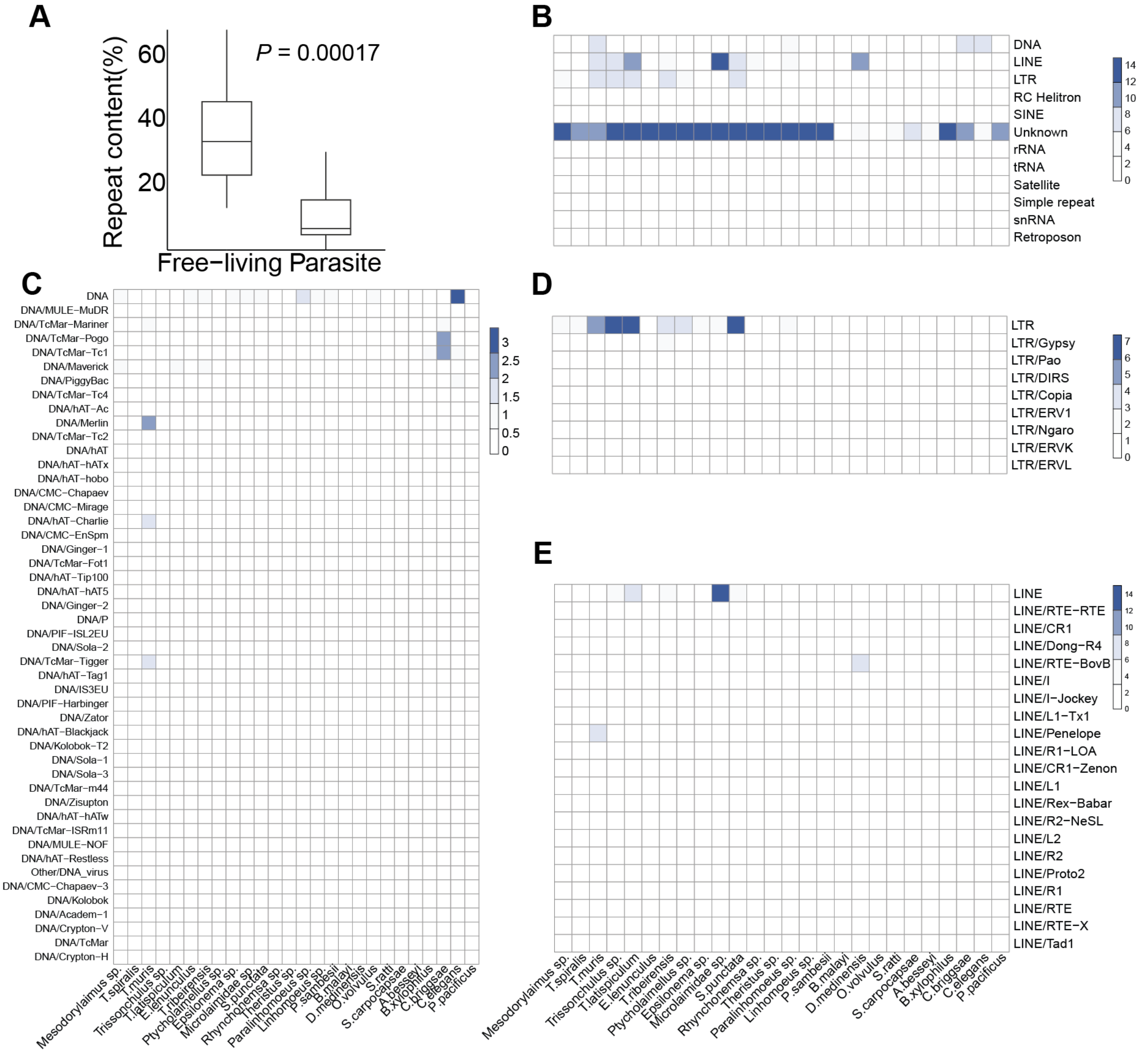

**Figure S11. The Nematoda species tree.** The phylogeny tree was constructed with the proteomes of 26 nematodes with a available genome, which included 13 published repersentive species and 13 assemblies (bold font) in this study.The *P. caudatus* is place as the outgroup. The bootstrap support value of each node is 100.

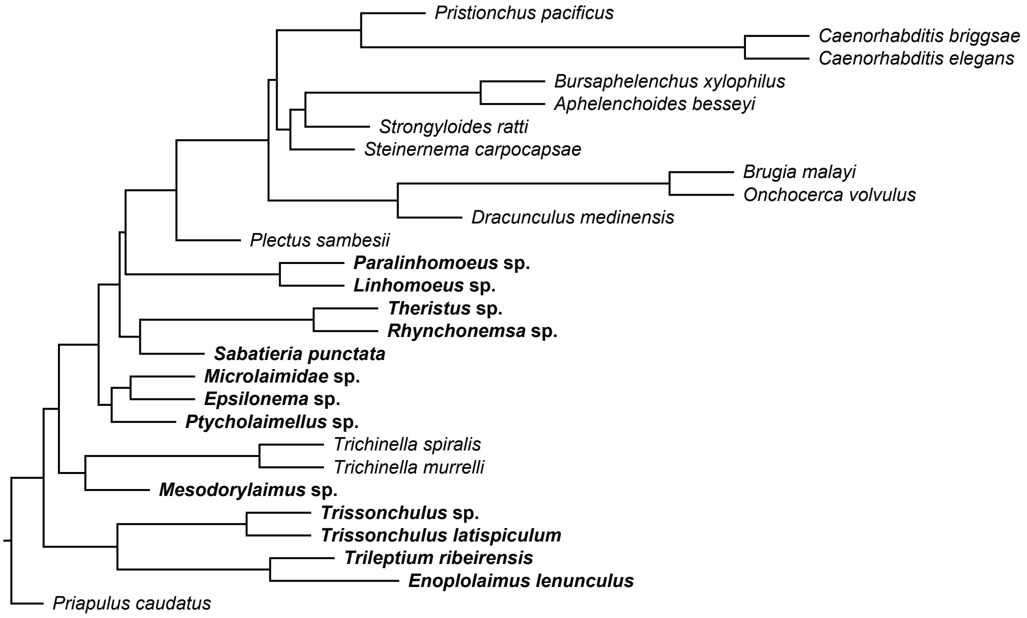
